# Germline Transmission of a Circular Human Artificial Chromosome in the Mouse

**DOI:** 10.1101/2022.06.22.496420

**Authors:** Aleksandra Wudzinska, Leslie A. Mitchell, Ran Brosh, Nazario Bosco, Francesco Boccalatte, Ioannis Aifantis, Sang Y. Kim, Teresa Davoli, Jef D. Boeke

## Abstract

Although the structure and function of the alphoid-tetO Human Artificial Chromosome (tetO-HAC) has been previously described in cell culture models and somatically in the mouse, *in vivo* persistence and stability throughout meiosis and across generations were not evaluated. Here we report germline transmission of a circular tetO-HAC across three mouse generations without observable health or reproductive deficiencies. Furthermore, we show that the tetO-HAC is maintained without selection as an episome and can be efficiently transmitted by both ova and sperm.

## Introduction

Mammalian artificial chromosomes (MACs) are a promising gene delivery approach offering the potential to add hundreds of new genes to cells complete with native regulatory elements without altering endogenous genomes. This capability offers a key advantage over traditional viral-based gene delivery techniques^1–3^, such as retroviral/lentiviral vectors, where payloads are limited and rely on genome integration. In addition, employing MACs may circumvent other shortcomings such as off-target insertional mutagenesis, immunogenicity and toxicity^4,5^.

The alphoid-tetO Human Artificial Chromosome (tetO-HAC) was previously constructed through a bottom-up approach based on a 348-bp alphoid dimer, where one monomer contains a 17-bp naturally occurring Centromere protein B (CENP-B) box allowing kinetochore assembly and the other monomer has this sequence swapped with a 42-bp tetracycline operator (tetO) binding motif, allowing conditional loss through kinetochore deactivation by binding of Tet Repressor (tetR) proteins to tetO repeats^6–10^. The dimer was amplified up to a ∼50-kb alphoid array, cloned into a blasticidin-resistant YAC/BAC that served as input DNA for multimerization in human fibrosarcoma HT1080 cells, producing the final 1.1-Mb circular tetO-HAC. Subsequently the tetO-HAC was moved twice by microcell-mediated chromosome transfer (MMCT), first to DT40 chicken cells for insertion of a loxP-5’-HPRT1-Hyg-TK cassette, and then to *HPRT1*-deficient Chinese Hamster Ovary (CHO) cells, where the site was loaded with eGFP-3’-HPRT1 reporter by Cre/lox-mediated recombination (**Fig. 1a**)^8^. The tetO-HAC seems to be transmitted as a true chromosome as evidenced by sister chromatids pairing and kinetochore assembly in cell culture^7,11^. Interestingly, during formation in HT1080 cells, the tetO-HAC also appears to have captured a ∼4.2-Mb gene-poor genomic fragment of chromosome 13^9,11^. Although the authors did not fully investigate the significance of this incorporation, the resultant ∼5-fold increase in size may have enhanced the tetO-HAC’s compatibility with mammalian cells that may have evolved to harbor only large chromosomes^12,13^.

**Figure 1.**
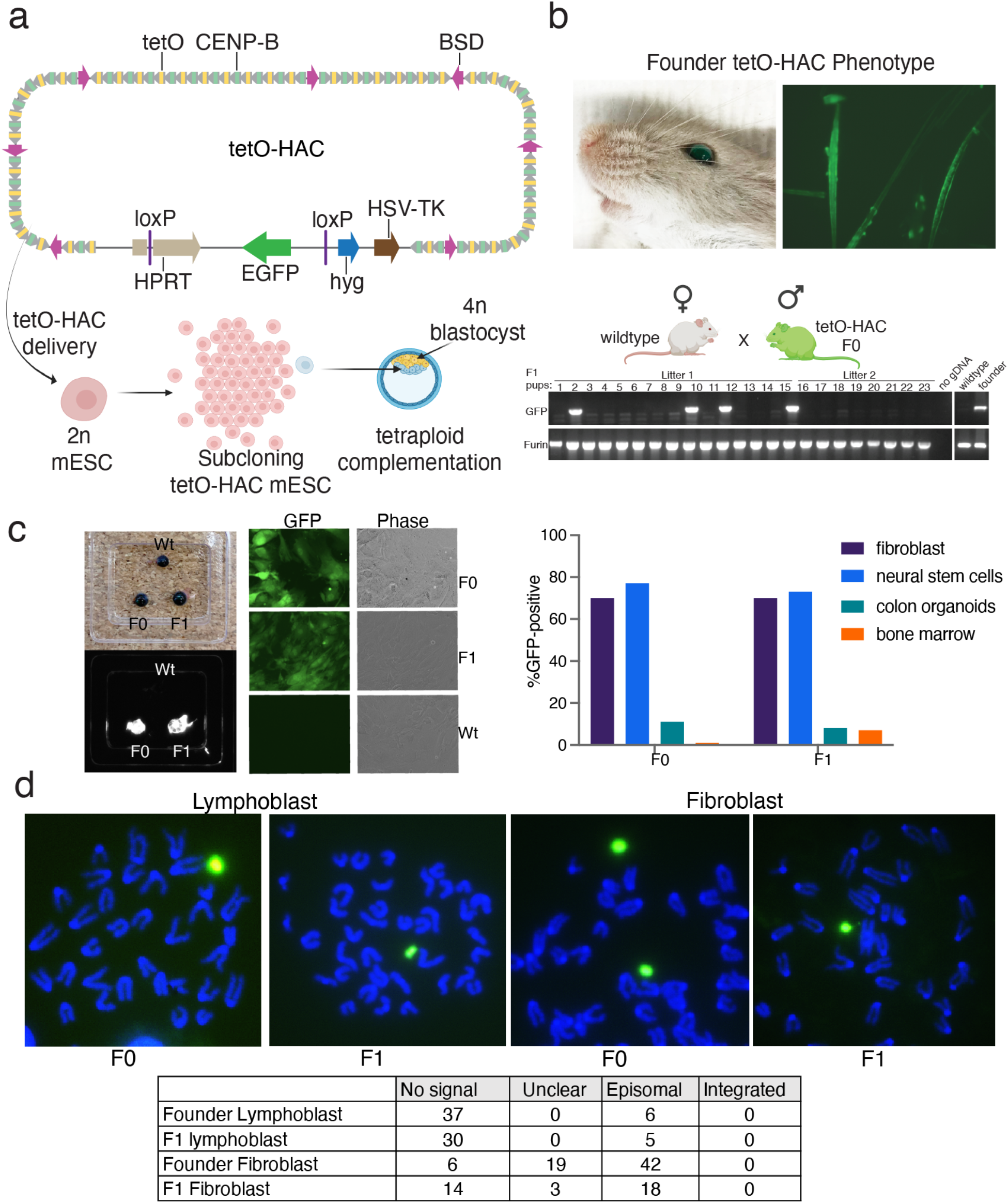
(**a**) Top, tetO-HAC structure and mouse derivation (illustrated with BioRender.com). The alphoid array contains multiple copies of blasticidin-resistance gene (BSD). The tetO-HAC also features a loxP-5’-HPRT1-Hyg-TK (Herpes Simplex Virus Thymidine Kinase) which was loaded with a 3’HPRT1-eGFP-hyg cassette (enhanced green fluorescent protein-hygromycin resistance gene (hyg)). (**b**) Top, F0 male with green-tinged eyes and green fluorescent tail hair. Bottom, genotyping F1 progeny of a tetO-HAC male crossed with a WT female. eGFP detection confirms tetO-HAC presence. Furin serves as loading control. (**c**) eGFP fluorescence images of whole dissected eyes (left) and cultured fibroblasts (right) from F0 and F1 progeny. Right, flow cytometry quantification of eGFP fluorescence for cultured fibroblasts, neural stem cells, colon organoids and bone marrow, derived from F0 and F1 mice. (**d**) DNA-FISH imaging of metaphase spreads prepared from primary fibroblasts and lymphoblasts harvested from F0 and F1 mice. Bottom, quantification of metaphase spreads, unclear denotes that chromosomes were not spaced to allow definitive determination of episomality or integration, but are included in the tally for transparency.

## Results

To accelerate the process of generating a tetO-HAC mouse in our laboratory and study its potential transmission, we obtained a previously reported^14^ mouse embryonic stem cell (mESC) line carrying the tetO-HAC that had been transferred from CHO cells through MMCT (**Fig. 1a)**. In our hands, the tetO-HAC mESC line exhibited considerable growth variability and clonal heterogeneity. Consequently, we applied hygromycin selection to enrich for the presence of the tetO-HAC and generated a series of sub-clones before attempting tetraploid complementation, as described previously^15–17^, a method that yields mice entirely derived from injected mESCs. We attempted 13 separate injections with different clones with almost all cases resulting in spontaneous abortion at ∼E16 or death of the pups shortly after birth. We did ultimately obtain a single live birth of a male mouse, which presented a green-tinged eye and green fluorescent hair by microscopic imaging (**Fig. 1b, top**) at maturity, suggesting the acquisition and maintenance of the tetO-HAC in at least some tissues, consistent with a previous report^14^.

To expand the colony and evaluate potential tetO-HAC germline transmission, we paired the founder male with C57BL/6J females but obtained no pups despite multiple attempts. Replacing the C57BL/6J females with the highly fecund CD-1 strain proved fruitful, yielding two F1 litters for a total of 9 females and 15 males (**Fig. 1b, bottom**). PCR genotyping of tail biopsies revealed that 4/15 males and 0/9 females carried the eGFP gene, representing to our knowledge the first reported instance of germline transmission of the tetO-HAC.

To determine the level of eGFP protein expression, we sacrificed the founder and an F1 male and collected multiple tissues for analysis, as well as sperm for future *in vitro* fertilization. Both tetO-HAC mice exhibited high fluorescence in eyes and in cultured primary fibroblasts (**Fig. 1c, left**). Flow cytometry analysis revealed a high proportion of eGFP-positive cells among fibroblasts and neural stem cells, but not in intestinal cells derived from dissociated colonic organoids or red blood cell depleted bone marrow (**Fig. 1c, right**). Further analysis by flow cytometry showed that there was an enrichment of eGFP-positive cells in the hematopoietic stem and progenitor fraction (**Supplementary fig. 2**).

A critical feature of this particular chromosome is its potential to be conditionally lost through inactivation of the kinetochore, which is only feasible if the tetO-HAC can replicate and segregate autonomously without genomic integration. To determine whether the tetO-HAC had integrated or remained autonomous, we prepared metaphase spreads of fibroblasts and lymphoblasts cultured from the F0 and F1 mice and detected the tetO-HAC by DNA fluorescence *in situ* hybridization (DNA-FISH). We found that the HAC, which was detected in about half of the metaphase spreads, was episomal in all instances where chromosomes were adequately spaced, with no instances of genomic integration observed (**Fig. 1d**). As in the case of eGFP protein expression, the proportion of cells that contained a tetO-HAC FISH signal varied between lymphoblasts and fibroblasts indicating a cell type specificity to maintaining the tetO-HAC *in vivo* (**Fig. 1d, lower**).

Thus far, all tetO-HAC-carrying progeny were male and had inherited the chromosome paternally. Because there are sex-specific differences during meiosis^18,19^ we wondered whether the tetO-HAC could be transmitted through ova and whether tetO-HAC-carrying females were viable. We performed *in vitro* fertilization using sperm collected from an F1 male and C57BL/6J eggs to generate the F2 progeny, which included, tetO-HAC-confirmed males and females. Both sexes were backcrossed to wild-type C57BL/6J mice and the resulting F3 litters were genotyped. Breeding tallies showed that the tetO-HAC was inherited at approximately 24% when the dam transmitted the tetO-HAC as compared to 31% when sired (**Fig. 2a**). Notably, the F2 generation were competent breeders, averaging approximately a litter per month at 4.3 pups per litter, which suggests that the issues with breeding experienced with the founder may have been a byproduct of tetraploid complementation rather than being tetO-HAC-related. The proportion of fibroblasts expressing eGFP remained high in the F3 progeny of both sexes (**Fig. 2b**). Importantly, the tetO-HAC continued to be transmitted solely as an episome after two additional cycles of meiosis (**Fig. 2c**).

**Figure 2.**
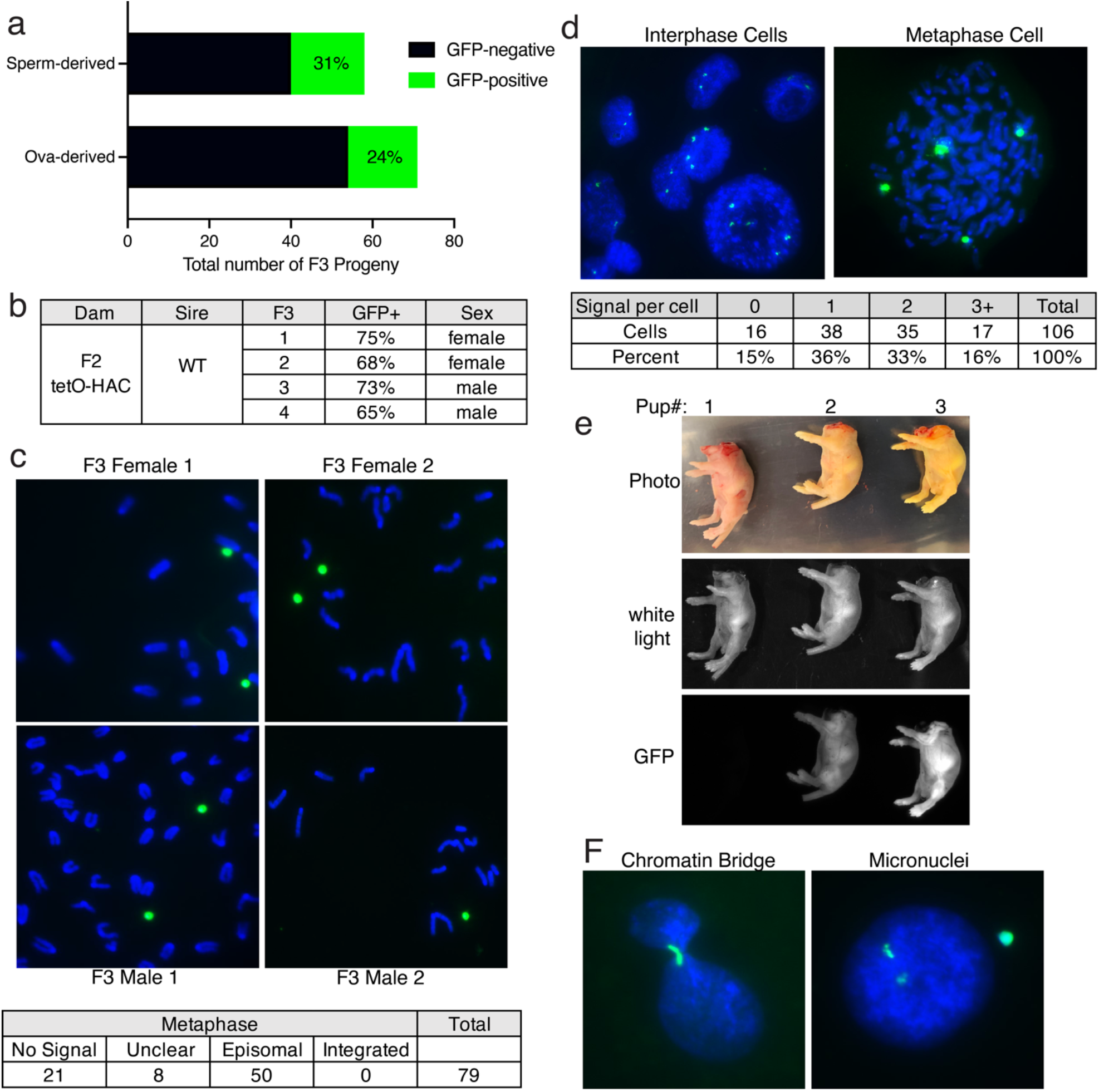
(**a**) Sperm- (male) and Ova- (female) derived tetO-HAC transmission rates. (**b**) Summary of eGFP fluorescence detection of Ova-derived tetO-HAC F3 progeny by flow cytometry of cultured fibroblasts. (**c**) Representative images and summary table of DNA FISH of metaphase spreads for F3-derived fibroblasts. (**d**) Representative images and summary table of interphase and metaphase cells exhibiting variable tetO-HAC copy number by DNA FISH analysis. (**e**) eGFP variability among F3 littermates. (**f**) DNA FISH of fibroblasts depicting a lagging tetO-HAC during telophase and tetO-HAC within micronuclei.

## Discussion

Although mitotically and meiotically stable without selection, we noted considerable variability in tetO-HAC ploidy when examining interphase cells where DNA-FISH signals ranged from 0 to 6, with roughly half of fibroblasts showing more than one signal (**Fig. 2d**). This heterogeneity was also observed for eGFP expression among individuals of the same litter (**Fig. 2e**). Although not verified, this may be a consequence of the number of tetO-HAC copies inherited from the germ cell at conception.

Why the tetO-HAC is present at variable percentages in different mitotic cell types is up to speculation, but the reason may simply be due to different cells’ abilities to tolerate substantial eGFP expression, as some studies have shown^20,21^, or perhaps some feature of the HAC’s structure makes it more often lost from certain populations of cells. Importantly, at the organismal level, the tetO-HAC does not seem to cause any obvious physiological phenotype and it is quite possible that when loaded with a less toxic, even therapeutic payload, these cell type-specific differences would ameliorate. We also showed that both males and females can efficiently transmit the tetO-HAC at approximately the same rate. Interestingly, in a previous study using a different circular HAC^22^, the transmission to offspring from females remained constant at 40% while that from males decreased from 50% to 0% with successive matings, underscoring that not all circular HACs behave the same in this respect.

However the aforementioned hypothesis does not explain why tetO-HAC copy number is so variable between cells of the same type. Segregation errors occur when a single kinetochore is tethered to both poles leading to chromosome lagging and subsequent aneuploidy ^23,24^. We occasionally see evidence of this type of misegregation in primary fibroblasts where mitotic cells are stalled at telophase with the tetO-HAC observed in a chromatin bridge, struggling to segregate to either pole (**Fig. 2f, left**) and corroborated by detection of micronuclei, a resolution to bridged chromatin resulting in genomic exclusion from either daughter cell^25^ (**Fig. 2f, right**). In fact, the cytokinetic defects caused by these bridges can lead to endoreduplication^26,27^, where genomic DNA is replicated in the absence of cytokinesis leading to polyploidy, as depicted in (**Fig. 2d, right**). This may partially explain the variation in tetO-HAC copy number discussed earlier.

Here we show that large multi-Mb circular chromosomes with human-specific alphoid DNA engineered to be conditionally unstable can be transmitted maternally and paternally through the germline in the mouse and be maintained as an episome without chemical selection not only throughout ontogeny but also across multiple generations and through both germ lines.

## Acknowledgements

We thank Mikhail Liskovykh for providing the clone B1 tetO-HAC mESC line. We thank Nicholas Lee for preliminary work contributing toward this study, and for helpful discussions on HAC molecular and cellular biology. This work was supported in part by DARPA contract HR0011-16-2-0002 and NIH/NHGRI grant 1RM1HG009491 to JDB.

## Methods

### Tetraploid complementation

Tetraploid complementation injections were performed as previously described ^17,28,15,16^. Briefly, female B6D2F1 mice were superovulated by intraperitoneal injection with pregnant mare serum (PMS) and human chorionic gonadotropin (hCG) and crossed to B6D2F1 stud males. Fertilized zygotes were isolated from females that had a visible vaginal plug 24 h after hCG injection. Zygotes were cultured until the 2-cell stage and were electrofused. After 1 h, 4n embryos were carefully separated, cultured in KSOM (Simplex Optimized Medium with Potassium) medium for another 2 days, and then injected with 15–20 mESCs before embryo transfer to pseudopregnant CD-1 females. The NYU Langone Health Animal Care and Use Committee approved all procedures described in this work.

### Genotyping

Ear punches were collected from weaned mice, serving as a unique identifier among cage mates. Genomic DNA was obtained by lysing ear punches in DirectPCR Ear Lysis Reagent (Viagen, 102-T) according to the manufacturer’s recommendation. For genotyping PCR, 1 µl of crude DNA lysate was used with 2x GoTaq Reaction Mix (Promega M7123) supplemented with 5% DMSO.

### Tissue Harvest and Flow cytometry

Fibroblasts and neural stem cells were collected and cultured as described^29,30^. For colon cells, the entire colon was harvested, set on bibulous paper and opened lengthwise. After thorough washing with PBS, the opened colon was incubated in 10 ml of 8 mM EDTA in PBS for 2 h at 4 °C and then shaken vigorously to separate colonic crypts. Crypts were plated in Matrigel (Corning 356235, 100-500 crypts per 50 ul per well in 24-well plate) and incubated for 20 min at 37 °C before adding 500 ul of complete colon organoid media (Supplementary Table 1). Resulting organoids were dissociated and eGFP was quantified by flow cytometry using live cells.

### Flow cytometry of bone marrow cells

Mouse bone marrow was flushed out from the long bones of the lower limbs of sacrificed animals. Single-cell suspensions were derived by mechanical disruption of bone marrow in PBS supplemented with 2% Fetal Calf Serum (FCS, Sigma, C8056). For FACS analysis, red blood cells were lysed for 5 min with BD PharmLyse (BD biosciences, 555899) and immediately washed. Nonspecific antibody binding was blocked by incubation with 20 µg/ml Rat IgG (Sigma, I4131) for 5 min. Cells were incubated with primary antibodies for 1-3 h. Antibodies used to detect hematopoietic stem and progenitors by flow cytometry are listed below:

| **Antigen** | **Conjugate** | **Clone** | **Dilution** | **Manufacturer** | **Cat #** | **Lot #** |
| --- | --- | --- | --- | --- | --- | --- |
| **Ly-6A/E (Sca-1)** | PE-Cy7 | D7 | 1:400 | Biolegend | 108114 | B260282 |
| **CD117 (c-Kit)** | APC | 2B8 | 1:400 | Biolegend | 105812 | B287105 |

Stained cells were quantified using a BD Fortessa analyzer (BD biosciences). Flowjo software (BD Biosciences) was used to generate flow cytometry plot and calculate fluorescence intensities.

For enrichment of hematopoietic stem and progenitor cells (HSPCs), whole bone marrow was processed by magnetic selection with anti-CD117 microbeads (Miltenyi Biotec, 130-091-332).

Bone marrow cells destined for metaphase analysis, were enriched for HSPC and cultured in OptiMEM (Gibco, 31985-062) supplemented with 15% FCS (Sigma, C8056), 100 U/ml penicillin-streptomycin (Cellgro, 30-002-Cl), 100 µg/ml Normocin (Invivogen, ant-nr-1), and mouse recombinant cytokines (100 ng/ml SCF (Peprotech, 250-03), 50 ng/ml Flt3-L (Peprotech, 250-31L), 10 ng/ml IL-6 (Peprotech, 216-16), 10 ng/ml IL-3 (Peprotech, 213-13) and 10 ng/ml TPO (Peprotech, 315-14)). Cells were cultured for 48h before proceeding with metaphase analyses.

### Metaphase spreads

Cells at 70% confluence were arrested at metaphase with colcemid (Sigma, 10295892001, 0.2 µg/ml, 4 h) collected by trypsinization and resuspended in hypotonic 0.075 M KCl solution for 10 min at room temperature. Cells were pre-fixed by adding 1 ml of cold fixative (methanol/acetic acid 3:1) for 5 min, resuspended drop-wise in pure fixative and dropped onto cold slides.

### DNA-FISH

The fluorescent probe was prepared using a 50 kb BAC containing the tetO-HAC alphoid array (gift from Vladimir Larionov) and labeled with Biotin-16-dUTP via nick translation as described previously^31^. Slides were pre-treated with RNAse A (EN0531, 100 µg/ml, 1 hr at 37 °C), dehydrated in an ethanol gradient (70%, 90%, 100%, 5 min each) and denatured (25% formamide, 4X SSC at 85 °C, 3 min). Probes were hybridized to metaphase spreads (200 ng of prepared probe, 4X SSC, 10% dextran sulfate, 25% deionized-formamide) overnight at 37 °C in a moist chamber. The hybridized biotin-labeled probe was detected with FITC-conjugated Avidin for 30 min before mounting with VECTASHIELD medium containing DAPI (Vector Laboratories, H-1200) and imaged using EVOS M7000 imaging system (Thermo Fisher, AMF7000).

