## Supplementary Figure 1 for "Germline Transmission of a Circular Human Artificial Chromosome in the Mouse"

### Slide 1
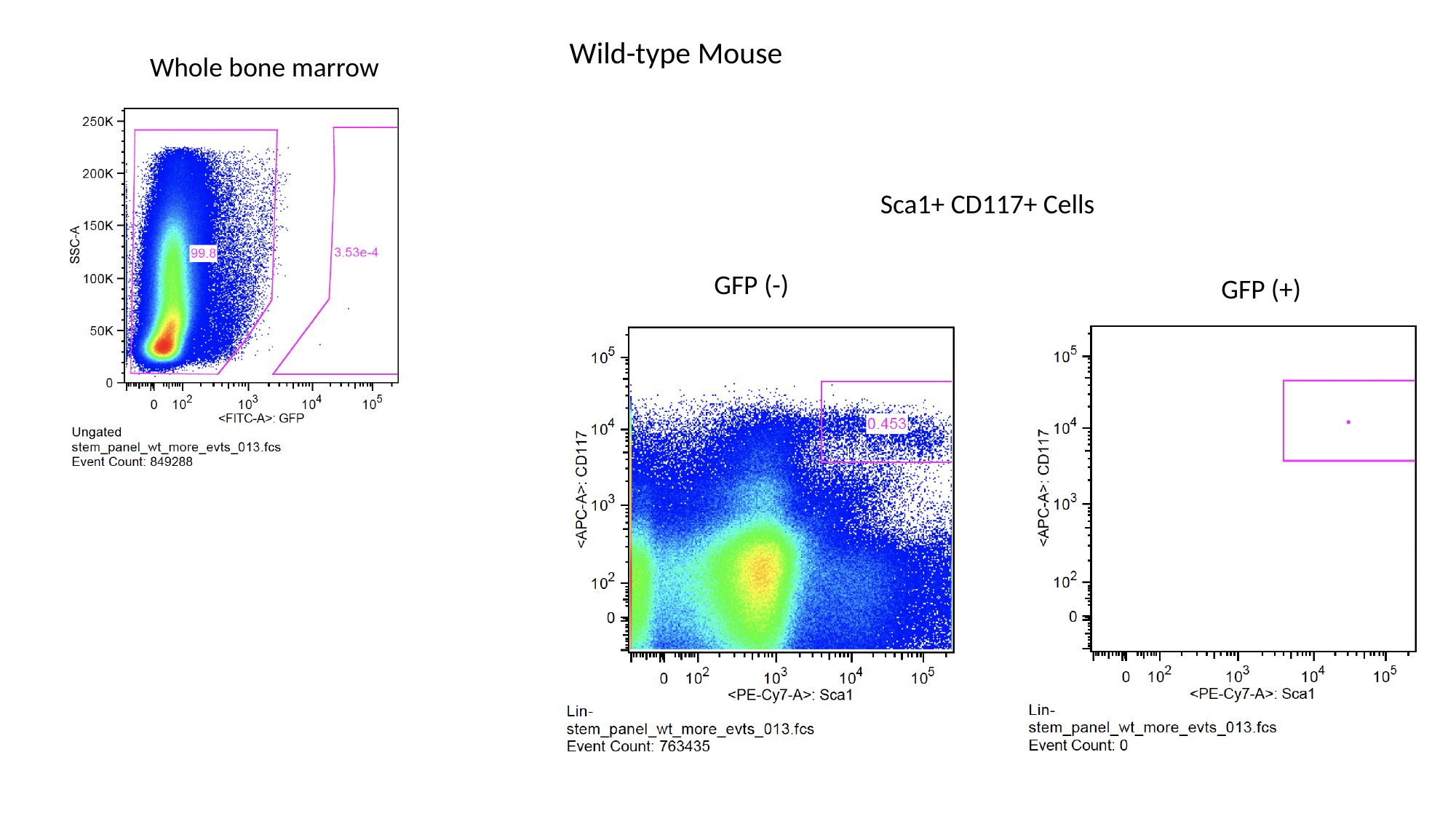

Wild-type Mouse
Whole bone marrow
Sca1+ CD117+ Cells
GFP (-)
GFP (+)

### Slide 2
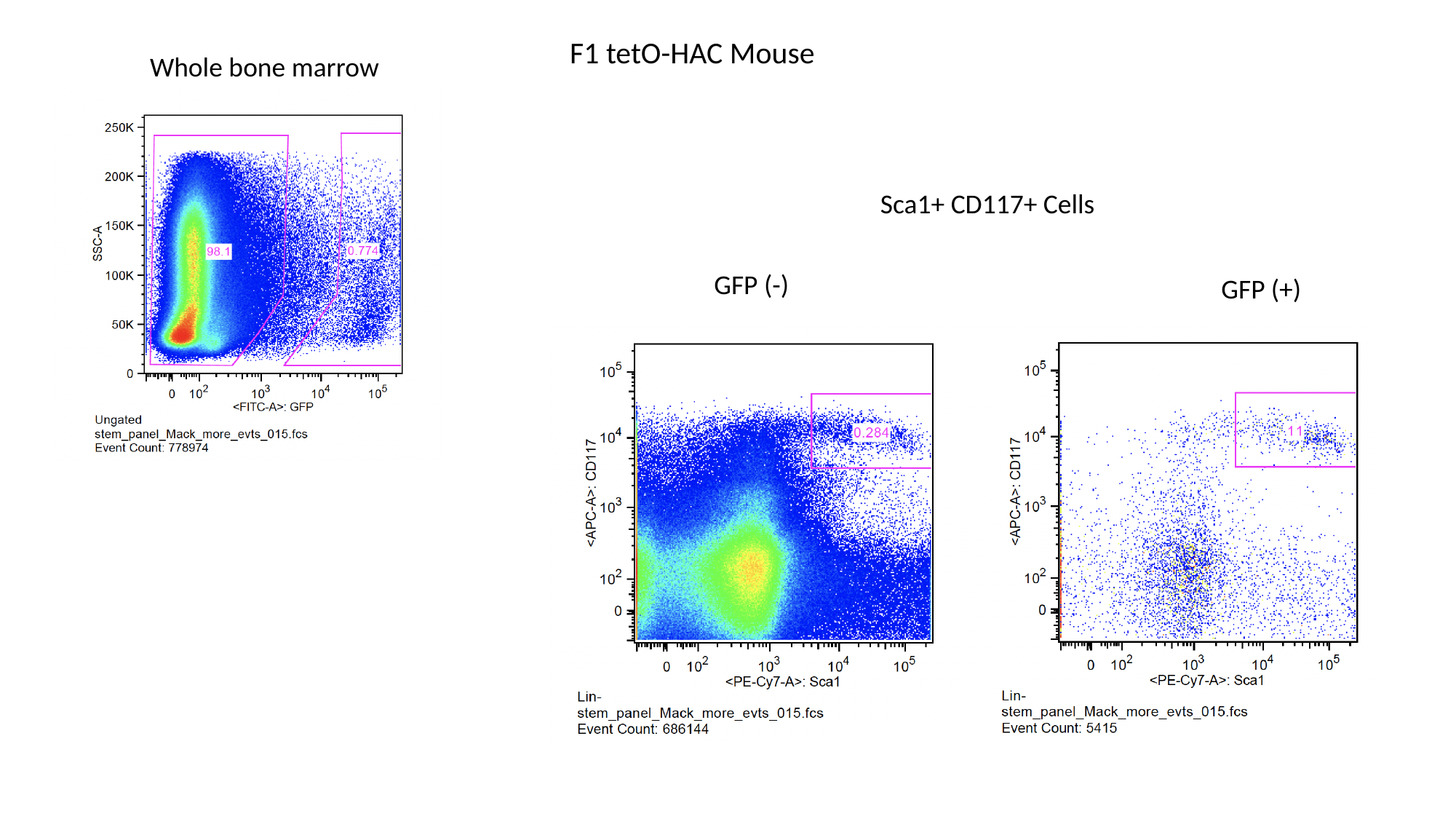

F1 tetO-HAC Mouse
Whole bone marrow
Sca1+ CD117+ Cells
GFP (-)
GFP (+)

### Slide 3
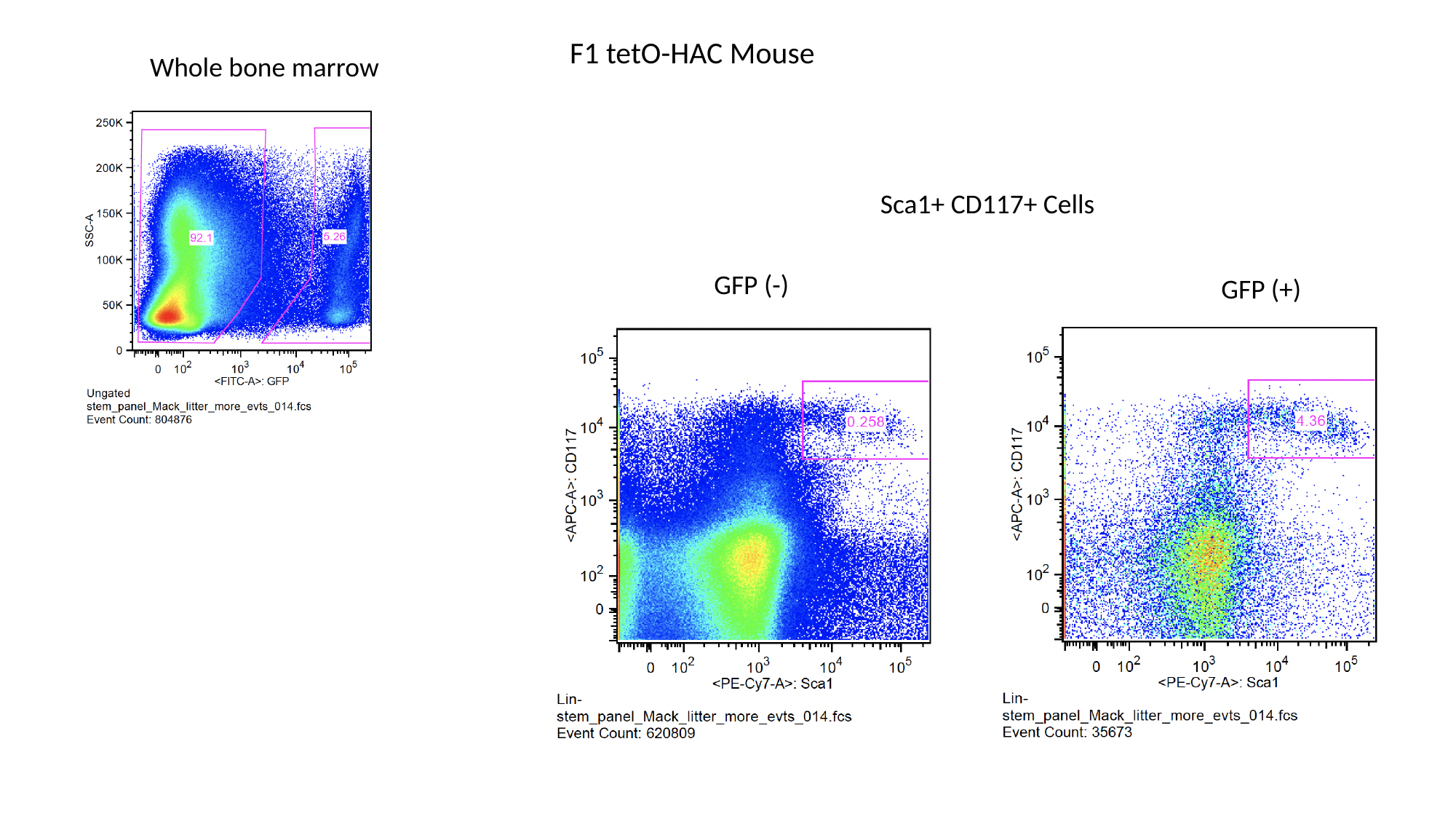

F1 tetO-HAC Mouse
Whole bone marrow
Sca1+ CD117+ Cells
GFP (-)
GFP (+)
