## Supplementary Table 1 for "Germline Transmission of a Circular Human Artificial Chromosome in the Mouse"

| ccm (mouse colon organoids) | | | |
| --- | --- | --- | --- |
| **Component** | **Stock Con.** | **Final Con.** | **Volume**  **(in 50ml)** |
| Advanced DMEM/F-12 Medium  (Gibco 12634028) | - | - | 47.5ml |
| B27 supplement  (Gibco 17504044, Store at -20ºC) | X50 | X1 | 1ml |
| GlutaMAX supplement  (Gibco 35050061, Store at RT) | X100 | X1 | 500ul |
| HEPES  (Gibco 15630080, Store at 4ºC) | 1M | 10mM | 500ul |
| Pen/strep | X100 | X1 | 500ul |
| N-2 supplement  (Gibco 17502048, Store at -20ºC) | X100 | X1 | 500ul |
| N-Acetylcysteine  (Sigma Aldrich A9165-5G, Dissolved in 3’D.W, Store at 4ºC) | 500mM | 1.25mM | 125ul |
| Recombinant Human EGF  (Gibco PHG0311, Dissolved in 1% BSA/PBS, Store at -20ºC) | 50ng/ul | 25ng/ml | 25ul |
| Noggin  (Peprotech 250-38, Dissolved in 1% BSA/PBS, Store at -20ºC) | 100ng/ul | 100ng/ml | 50ul |
| R-Spondin-1  (Peprotech, 120-38) | 1ug/ul | 1ng/ml | 50ul |
